# A deep learning approach to predict the impact of non-coding sequence variants on 3D chromatin structure

**DOI:** 10.1101/516849

**Authors:** Tuan Trieu, Ekta Khurana

## Abstract

Three-dimensional structures of the genome play an important role in regulating the expression of genes. Non-coding variants have been shown to alter 3D genome structures to activate oncogenes in cancer. However, there is currently no method to predict the effect of DNA variants on 3D structures. We propose a deep learning method, DeepMILO, to learn DNA sequence features of CTCF/cohesin-mediated loops and to predict the effect of variants on these loops. DeepMILO consists of a convolutional and a recurrent neural network, and it can learn features beyond the presence of CTCF motifs and their orientations. Application of DeepMILO on a cohort of 241 malignant lymphoma patients with whole-genome sequences revealed CTCF/cohesin-mediated loops disrupted in multiple patients. These disrupted loops contain known cancer driver genes and novel genes. Our results show mutations at loop boundaries are associated with upregulation of the cancer driver gene *BCL2* and may point to a possible new mechanism for its dysregulation via alteration of 3D loop structures.

## 1. Introduction

Human genome is organized into three-dimensional (3D) hierarchical structures such as chromosomal compartments, topologically associated domains (TADs), and chromatin loops. Chromosome conformation capture techniques such as Hi-C ^1^ and ChIA-PET ^2^ can be used to identify these 3D structures. In particular, ChIA-PET assays capture chromatin interactions between loci mediated by a specific protein factor ^2 3^. Multiple studies have found that DNA loops mediated by CTCF and cohesin (SMC1, SMC3, RAD21 and either STAG2 or STAG1) bound on both anchors at the loop ends isolate genes from active enhancers and their disruption can cause dysregulation of nearby genes ^4 5 6 3 7^. Mutations in boundaries of such insulator loops may break or weaken loops and allow proto-oncogenes to interact with enhancers outside of the loops or to inhibit regulatory elements of tumor suppressors from interacting with their proper target genes ^5^. Thus, methods to identify the mutations that are likely to disrupt the chromatin loops are needed.

To the best of our knowledge, there is currently no method to identify mutations that can alter insulator loops. A natural approach is to model loops and then observe how loops are changed in the presence of mutations. However, modelling loops is challenging because the precise DNA sequence rules and mechanism of loop formation are not clear. While the majority of CTCF/cohesin-mediated loops are ‘hairpin loops’ ^3^ with boundaries containing CTCF motifs in convergent orientation, boundaries with tandem CTCF motifs (i.e. CTCF motifs with the same orientation) can form ‘coiled loops’ ^3^. Moreover, multiple studies have shown that transcription factors (TFs) other than CTCF and cohesin may play an important role in loop formation ^89^. Recent experiments ^7^ support the loop extrusion model ^10 11^, which suggests that structural maintenance of chromosome (SMC) proteins (e.g. cohesin or condensin) extrude chromatin until blocked by two CTCF proteins bound at convergent CTCF motif sites to form loops. Yet, it is not clear how and when CTCF proteins can prevent SMC proteins from extruding, and how SMC proteins can translocate along the chromatin at the rapid speeds observed in experiments ^7 12 13^. In modelling insulator loops, a model has to learn DNA sequence patterns of CTCF-bound regions that can stop cohesin proteins from extruding because most CTCF-bound regions in fact do not form loops ^14^. It becomes even more challenging to model loops when DNA sequence of anchor regions contains multiple CTCF motifs with opposite orientations. Additionally, loops involve two loci, a start point and a stop point of the extrusion process, and loops can interconnect ^3^ so the model should be able to identify pairs of DNA fragments of regions that form loops. Recently, Hansen et al. ^15^ found evidence of RNA binding CTCF proteins to mediate a class of chromatin loops that show different DNA sequence patterns compared with RNA-independent loops. Thus, it is clear that there are more DNA sequence patterns besides CTCF motifs that are important for loop formation and methods relying solely on the presence of CTCF motifs and/or CTCF motif orientation to predict or to model loops are unlikely to do well.

Lollipop ^16^ is a computational method that attempts to predict CTCF-mediated loops from a range of genetic and epigenetic features. However, the method cannot be used to predict the impact of mutations on loops because it does not take into consideration specific DNA sequences and therefore, it cannot account for sequence differences caused by mutations. CTCF-MP ^17^ also predicts CTCF-mediated loops from genetic and epigenetic features. It uses a model based on word2vec ^18^ to learn DNA sequence features and boosted trees to predict loops. CTCF-MP can account for sequence changes by mutations and can be used to predict the impact of mutations on loops. In spite of that, the word2vec model may not be able to learn complex DNA sequence features as evidenced by the inability of CTCF-MP to deal with loop boundaries containing several CTCF motifs.

Convolutional neural network (CNN), a class of deep learning neural networks, has been successfully used to learn DNA sequence patterns such as those for DNA and RNA binding proteins ^19^, DNA methylation ^20^ or chromatin-profiling data ^21^. Another class of deep neural networks is recurrent neural network (RNN), which is commonly used for learning tasks involving sequential data such as language translation and speech recognition. Yet, it has not been used widely on DNA sequence, which is a type of sequential data where the order and relationship between the bases are important for its function. While CNNs are good at capturing local patterns in sequences, RNNs like long short-term memory (LSTM) networks can capture long distance dependencies in sequential data. Here, we show that RNN can perform comparably with CNN model in learning DNA sequence patterns of boundaries of insulator loops. Furthermore, they learned different features and combining their features delivered a better model compared to individual RNN or CNN models. Using features learned by a CNN and an RNN, we propose DeepMILO, a **Deep** learning approach for **M**odeling **I**nsulator **LO**ops, to learn DNA sequence features of insulator loops. The model can separate DNA sequences of insulator loop boundaries bound by both CTCF and cohesin proteins from DNA sequences of CTCF ChIP-seq peaks bound by only CTCF and DNA sequences of regions without CTCF binding. DeepMILO can pair DNA sequences of boundaries forming loops to distinguish insulator loops from different types of non-loops (i.e. fake loops) with high accuracy. Using DeepMILO, users can predict the impact of variants obtained by whole-genome sequencing of their samples on insulator loops from the cell-type of interest (Figure 1). Application of DeepMILO on 241 samples of malignant lymphoma patients from International Cancer Genome Consortium (ICGC) identified insulator loops changed by mutations in multiple patients. Our results reveal novel genes and known driver cancer genes whose expression may be altered in tumors via alteration of 3D chromatin structures.

**Figure 1.**
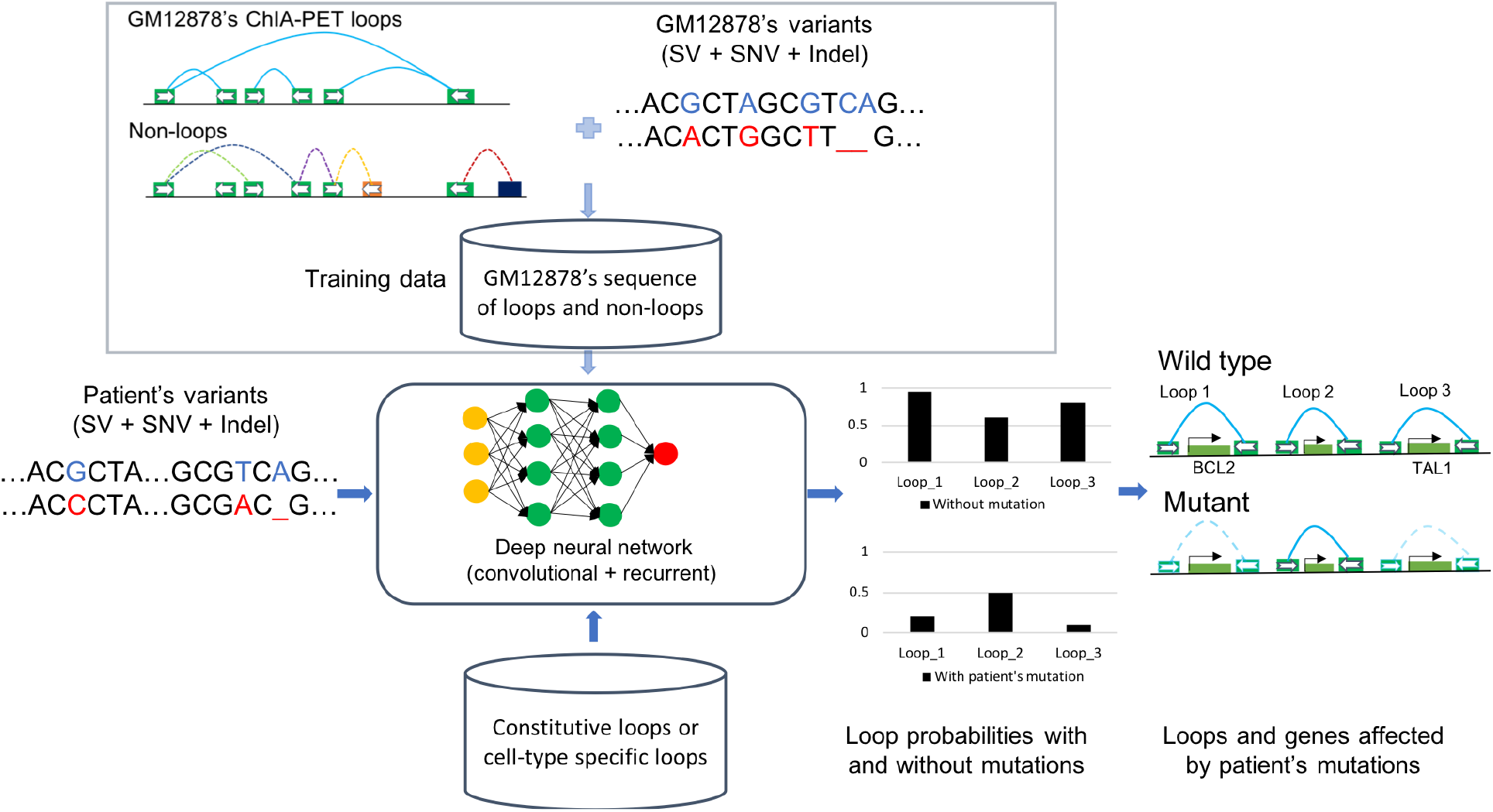
Overall approach for identification of affected insulator loops and genes by mutations in patients. Five different types of non-loops for model training are colored in different colors. Mutated bases in sequences are highlighted in red and blue. Boundaries of insulator loops and non-loops are used to train a model to learn sequence features of insulator loops. The model is then applied to patients, mutation data to detect insulator loops changed by mutations (boundaries of disrupted loops are connected by dotted lines)

## 2. Results

We first developed a ‘boundary model’ to learn DNA sequence features of boundaries or anchor regions of insulator loops. To pair two boundaries to form a loop, we built a model to distinguish left vs. right boundaries of loops using the learned features from the boundary model. This model is referred to as the ‘boundary orientation model’. The boundary and boundary orientation models were then combined to create DeepMILO with the capability of identifying boundaries and pairing them with their partners for loop prediction. Given a pair of DNA sequences of two boundaries, this model outputs a number between 0 and 1 that can be interpreted as the loop probability or loop strength. The models were trained, validated and tested with insulator loops of the cell line GM12878 captured by cohesin (RAD21) ChIA-PET with PET peaks overlapping CTCF ChIP-seq peaks ^5^. In particular, the data for chromosomes 7 and 8 was held out for testing and the data for chromosome 16 was used for validation. Additionally, the models were also tested with the data from chromosomes 7 and 8 of the cell line K562 ^5^. To increase the sensitivity of the models with mutations from patients, DNA sequences of boundaries were mutated with mutations from corresponding cell lines GM12878 ^22^ and K562 ^23^ for training, validating and testing. All datasets have approximately the same numbers of positive and negative samples and we used area under the receiver operating characteristic curve (AUROC) to measure the performance of the models.

### RNN can perform comparably with CNN and their features complement each other

We built a CNN model and an RNN model to extract features directly from DNA sequences of boundaries. The RNN model uses bidirectional long short-term memory blocks ^24^ (see **Methods**). The models were trained separately to identify boundaries of insulator loops. Individually, the RNN and CNN models show similar performance (AUROC of 0.744 for RNN and 0.746 for CNN) (Figure 2c) for the task of separating sequences of insulator boundaries (true boundary - Figure 2a) from sequences of CTCF ChIP-seq peaks (non-boundary type 1 - Figure 2a). Both types of sequences have active CTCF motifs but insulator boundary sequences are also bound by cohesin proteins. High AUROC values indicates that additional sequence rules govern the presence of insulator loop boundaries besides the presence of actively bound CTCF motifs in a given cell-type and our models are able to learn these rules. Moreover, this result demonstrates that the RNN is able to learn DNA sequence features and performs comparably with the CNN model.

**Figure 2.**
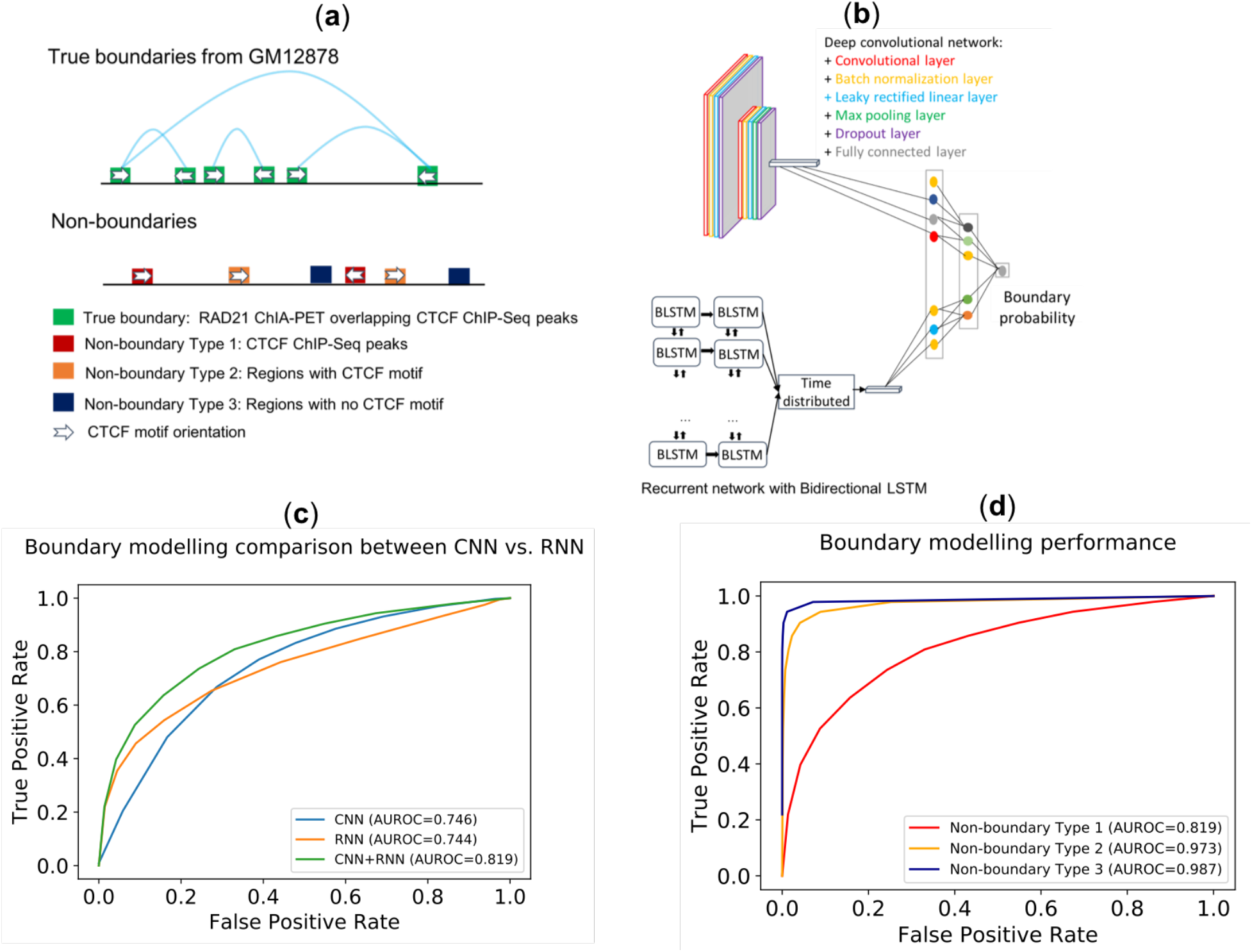
(**a**) Different types of non-boundaries used for training and testing. (**b**) Architecture of the boundary model consisting of a CNN and an RNN. (**c**) The RNN model performed comparably with the CNN model and their combination delivered the best performance. (**d**) Performance of the boundary model (CNN + RNN) on test sets from GM12878.

We then built the boundary model by combining learned features from the RNN and CNN models (Figure 2b). This model performed significantly better compared with individual RNN and CNN models for the non-boundary type 1 test set (AUROC of 0.819) (Figure 2c). The result suggests that the RNN and CNN models learned different sets of features that complement each other for the task of identifying DNA sequences of insulator loop boundaries. Again, the high AUROC demonstrates the capability of the model in learning features beyond actively bound CTCF motifs.

### Performance of the boundary model on different test sets

The boundary model was built from learned features of a CNN and an RNN as discussed above (Figure 2b). The model was tested with test sets containing boundaries from chromosome 7 and 8 of the cell line GM12878. We used three different test sets. They have the same set of true boundaries as positive samples but differ in negative samples (Figure 2a). The first test set contains negative samples from regions containing CTCF motifs overlapping CTCF ChIP-seq peaks of GM12878 but are not boundaries of any insulator loop (non-boundary type 1). The second set consists of regions with CTCF motifs but not bound by CTCF protein in GM12878 as negative samples (non-boundary type 2). The third test set includes regions without CTCF motifs and peaks as negative samples (non-boundary type 3). The first test set is the most difficult one as many negative samples can have similar DNA sequence patterns with positive samples.

The results from the three test sets are shown in Figure 2d. While the model performed well for the non-boundary type 1 test set achieving AUROC of 0.819 (also discussed above as part of Figure 2c CNN+RNN), it performed even better for other test sets. For non-boundary type 2, i.e. when negative samples contain just CTCF motifs but are not bound by CTCF protein, AUROC is 0.973. And as expected, the model performed best for non-boundary type 3, i.e. when negative samples contain no CTCF motifs or peaks (AUROC = 0.987). These results suggest that there are other sequence features rather than the presence of CTCF motif that distinguish boundaries of insulator loops and our model learned these features well.

For validation on an independent cell line, we created three similar test sets from chromosomes 7 and 8 of the cell line K562. The boundary model performed well with AUROCs of 0.78, 0.956 and 0.974 for test sets of non-boundary type 1, 2 and 3 respectively (see supplementary Figure S 1a). Thus, overall the model has learned sequence features of boundaries of insulator loops and generalized well as it was trained on data from GM12878 but also performed well on data from K562.

### Boundary orientation model to distinguish left and right boundaries

While most chromatin loops contain CTCF motifs in convergent orientation at the paired anchor regions, a boundary element can contain several CTCF motifs in different orientations. Moreover, the paired boundaries of some loops contain tandem CTCF motifs ^3^. Therefore, CTCF motif orientation alone cannot be used to distinguish the two boundaries of all loops. We used the learned features of the boundary model to build the boundary orientation model to distinguish left and right boundaries. This model shares its features with the CNN and RNN of the boundary model. Similar to the boundary model, the boundary orientation model was trained and tested with DNA sequences from the cell line GM12878. All boundaries in chromosomes 7 and 8 were used to test the model regardless of the number and orientation of their CTCF motifs. The results show that the left and right boundaries can be well separated with AUROCs of 0.97 and 0.957 for test sets from GM12878 and K562 respectively. We note that 49.4% of boundaries in the test set of GM12878 contain multiple motifs, which cannot be handled by methods relying on CTCF motif orientation. While models like CTCF-MP ^17^ or Lollipop ^16^ need to be fed with CTCF motif orientation explicitly, our results show that the boundary orientation model has learned features to distinguish left and right boundaries of loops de novo from the sequence.

### Loop model for learning features of pairs of DNA sequences of boundaries of insulator loops

As loops often interconnect and one boundary can be involved in several loops ^3^, pairing boundaries forming loops is not trivial. We combined the boundary model and boundary orientation model to build DeepMILO to predict insulator loops and effects of mutations on these loops (Figure 3b). The model was trained with data from the cell line GM12878 and tested with data from two cell lines GM12878 and K562.

**Figure 3.**
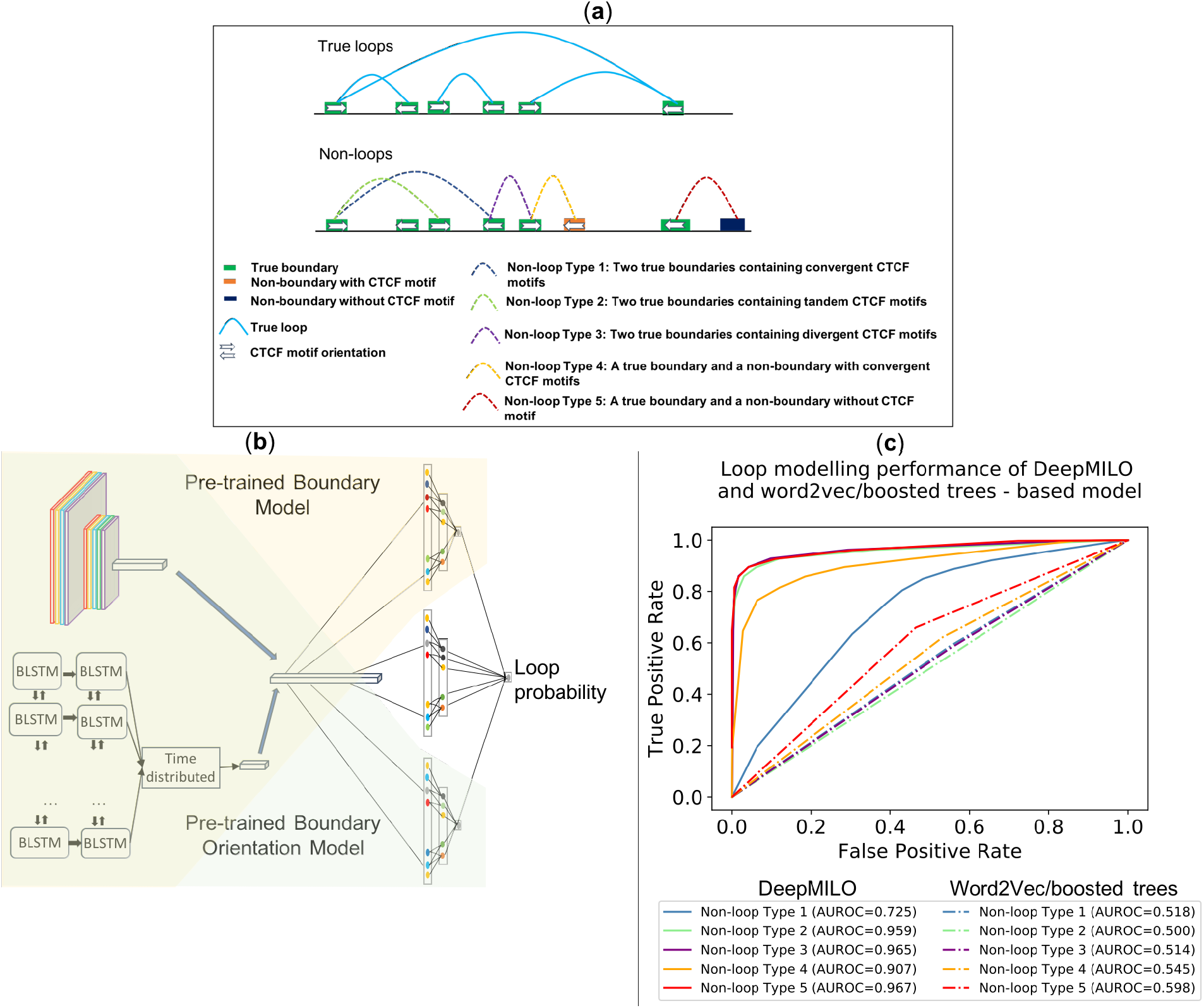
(**a**) Different types of non-loops for training and testing the loop model. (**b**) Details of DeepMILO; combining pretrained-boundary-model and pretrained-boundary-orientation-model helped training to converge faster. (**c**) Performance of DeepMILO and the model based on Word2Vec+Boosted Trees on test sets from GM12878

To ensure a fair evaluation of the model performance, five different types of test sets were created to test the model. They have the same set of positive samples from true insulator loops captured by CTCF/cohesin ChIA-PET. However, their negative samples are different as illustrated in Figure 3a. The first type of test set consists of non-loops formed from two true boundaries with convergent CTCF motifs (non-loop type 1). Each non-loop was derived from a true loop and shares one boundary with the true loop. The other paired boundary of the non-loop is the closest boundary to the paired boundary of the true loop and has a CTCF motif with the same orientation. This type of test set is the most difficult one as many non-loops possess similar patterns with true loops and they could be actual loops but were not captured in ChIA-PET experiments. The second type of test set includes non-loops from two true boundaries but containing tandem CTCF motifs (non-loop type 2). The third type of test set contains non-loops from two true boundaries with divergent CTCF motifs (non-loop type 3). The fourth type of test set consists of non-loops with one true boundary and one non-boundary containing a CTCF motif in convergent orientation with a CTCF motif at the true boundary (non-loop type 4). The fifth type of test set includes non-loops with one true boundary and one non-boundary containing no CTCF motif (non-loop type 5). The results from test sets of the cell line GM12878 are shown in Figure 3c. We also compared our deep learning model with a model based on word2vec ^18^ and boosted trees similar to CTCF-MP ^17^ but used DNA sequence features only.

For the most difficult test set, the non-loop type 1 test set, DeepMILO achieved AUROC of 0.725. A naïve model using CTCF orientation only would achieve an AUROC of ~ 0.5 as it would label all non-loops as loops. The word2vec based classifier achieved an AUROC of just 0.518 for the same test set. For the non-loop type 2 and 3 test sets, DeepMILO performed very well (AUROCs of 0.959 and 0.965 respectively). We note that the word2vec based classifier did not perform well on type 2 and 3 datasets with AUROC values of ~0.5 (Figure 3c). For the non-loop type 4 test set, DeepMILO also performed well with AUROC of 0.907. This test set contains non-loops with paired boundaries containing convergent CTCF motifs so that they are relatively difficult to classify. Models relying solely on CTCF motif orientation would achieve an AUROC of ~ 0.5 for this test set. The word2vec based classifier achieved an AUROC of 0.545 for this test set. Lastly, DeepMILO performed very well on non-loop type 5 test sets of non-loops with one non-boundary without CTCF motif (AUROC of 0.967). The word2vec based classifier performed better on the type 5 test set relative to the other sets achieving an AUROC of 0.598. Validation of DeepMILO on datasets of K562 shows similar results. DeepMILO achieved AUROCs of 0.691, 0.93, 0.937, 0.865 and 0.943 for test sets of non-loop type 1, 2, 3, 4 and 5 respectively (Figure S 1b), indicating that the model has generalized well.

DeepMILO performed significantly better than the model based on word2vec and boosted trees. A likely reason for the bad performance of the model based on word2vec and boosted trees could be the limitation of word2vec-based models in learning complex sequence features and in handling multiple CTCF motifs in boundaries of insulator loops. Insulator boundaries of CTCF/cohesin ChIA-PET loops are more complex than boundaries of CTCF-CTCF loops without cohesin binding and the word2vec-based model is not able to capture complex features in boundaries of insulator loops.

We provide DeepMILO as a software tool to predict the impact of variants on insulator loops. Given a set of variants, DeepMILO evaluates their impact on a set of ~ 14,000 constitutive loops shared by three cell lines GM12878, K562 and Jurkat ^5^ (default option) or a set of loops from the cell type of interest and outputs loop probabilities with and without variants (Figure 1). Users can then compare loop probabilities to identify altered loops. It is also possible to evaluate the effect of individual variants on insulator loops to identify functional variants.

### Validation of DeepMILO with known loop-disrupting deletions

We next tested DeepMILO on two known deletions that disrupt insulator loops. Hnisz et al. ^5^ introduced two deletions found in T-ALL patients using CRISPR/Cas9 at boundaries of loops containing the oncogenes *TAL1* and *LMO2*. The authors showed that the deletions increased interactions between enhancers and promoters that were insulated by boundary elements in the wild type cells leading to upregulation of *TAL1* and *LMO2* in the edited cells. We evaluated the impact of these two deletions on constitutive CTCF/cohesin loops. Among the ~14,000 loops, two loops cover the *TAL1* gene and three loops cover the *LMO2* gene (Figure S 2). Without any mutations, loop probabilities of the five loops calculated by DeepMILO are high in the range [0.80-0.84]. The deletion in neighborhood of *LMO2* is of length 25kb and affected 3 boundaries of three loops (Figure S 2). DeepMILO predicted loop probabilities of the three loops after the deletions are just ~ 0.06. The result indicates that the model correctly predicted the impact of the deletion on loop disruption. For the case of *TAL1*, the deletion is 400 bp (Figure S 2c). With the deletion, DeepMILO predicted the loop probabilities decreased to ~0.24, suggesting that the loops are seriously weakened or disrupted. These results are consistent with the experimental results in ^5^.

Given the large size of the two deletions, it is expected that they can disrupt the associated loops. To test the sensitivity of DeepMILO with small mutations and if small mutations are sufficient to disrupt insulator loops, we simulated 400 small deletions of one base for every position of the deletion related to *TAL1* and used DeepMILO to predict the impact of these small deletions. Comparing loop probabilities without and with individual mutations, we identified 11 consecutive mutations at the center of the deletion with highest reductions in loop probability (Figure S 2c). We find that the DNA sequence of these 11 positions matches well with a CTCFL motif (Figure S 2c). The reduction in loop probability is as high as 0.70, indicating that small mutations of one base could result in significant impact on insulator loops and that DeepMILO is sensitive to small mutations. These results demonstrate that DeepMILO can be used to predict the effect of variants from patients on insulator loops.

### Identification of disrupted loops in 241 malignant lymphoma patients and associated gene expression changes

The majority of somatic variants in cancer cells reside in non-coding regions ^25^. Some of these variants can affect 3D chromatin structures, which can in turn activate proto-oncogenes ^526^. We sought to apply DeepMILO to somatic variants from cancer patients to identify disrupted loops associated with non-coding variants. We selected the ICGC cohort with high number of single nucleotide variants (SNVs) (malignant lymphoma, MALY-DE) since it is much harder to predict the impact of smaller variants on loop disruptions relative to large genomic rearrangements that may delete or disrupt the entire loop regions. The cohort has 241 patients with whole-genome sequences and 104 patients also have matching RNA-seq data. We used the default ~14,000 constitutive insulator loops to identify loops changed in patients. When evaluating changes in loop probabilities, all mutations including SNVs, indels, inversions and copy number variants in boundaries of the loops were used although structural variants (SVs) spanning entire loops were excluded. Loop probabilities with and without mutations from patients were compared to identify loops that may be rewired to alternative conformations by somatic mutations.

Figure 4a shows the distribution of reductions in loop probability (defined as the difference between loop probability without mutations and loop probability with mutations). Most reductions are close to 0 (5^th^ and 95^th^ percentiles are −0.0024 and 0.0026 respectively), indicating that most loops do not change. This result is expected as most non-coding variants likely do not have major effects on loops. We then looked at the relationship between the number of mutated bases and the reduction in loop probability. The result shows that, as expected, more mutated bases in boundary elements is more likely to lead to larger changes in loop probabilities (Spearman’s rank-order correlation: 0.27). However, just a few mutated bases can cause large changes in loop probability (Figure 4c). These results are consistent with our result from the 400 small deletions of one base involving the *TAL1* gene’s loops.

**Figure 4.**
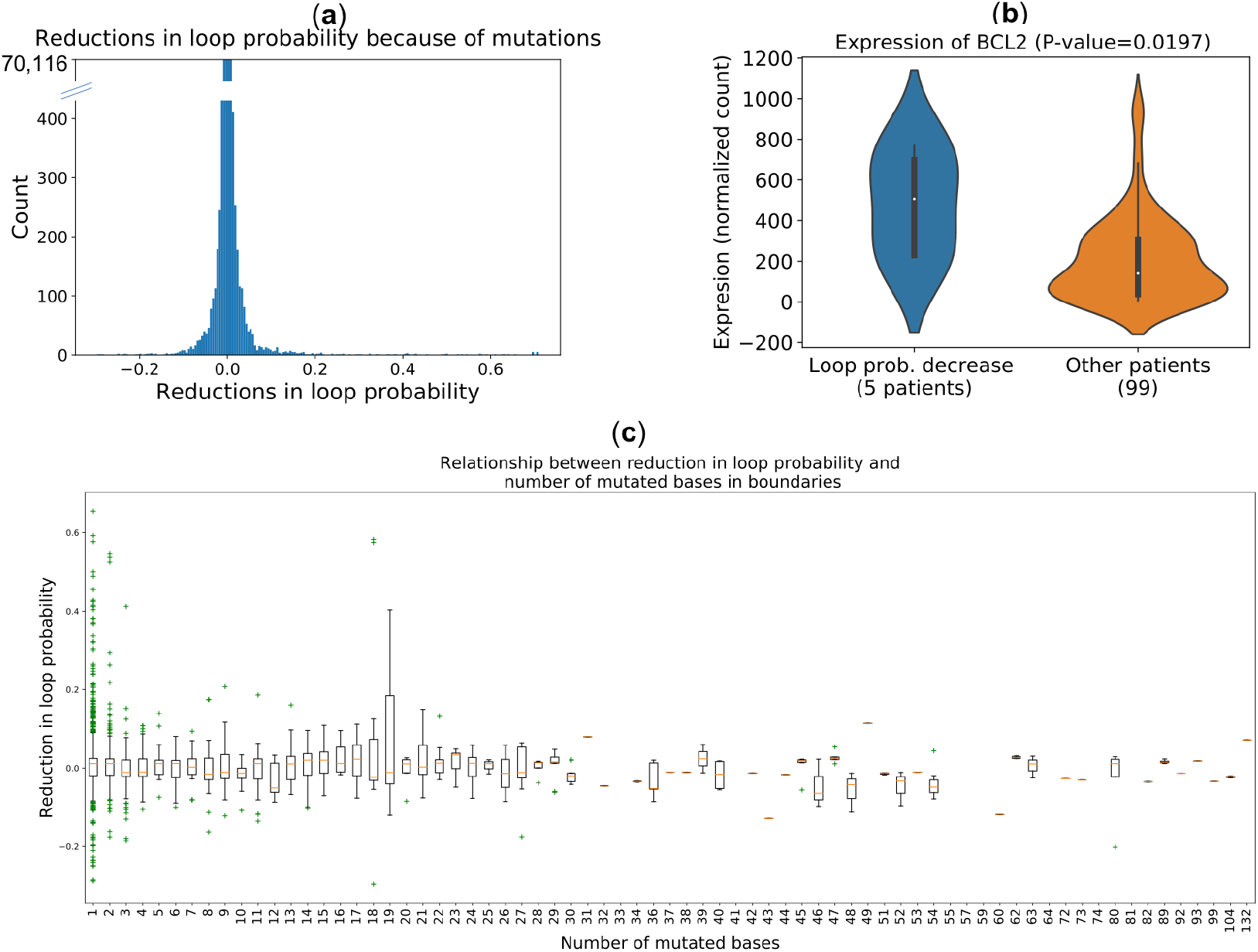
(**a**) Distribution of reductions in loop probability. (**b**) Oncogene BCL2 shows differential gene expression between patients with decrease in loop probability and other patients. (**c**) Relationship between reductions in loop probability with number of mutated bases in boundaries of loops in patients (loop probabilities with absolute values less than 0.01 were omitted for clearer trend; green crosses are outliers of box plots). Just few mutated bases can cause dramatic changes in loop probability.

We next analyzed the loops that are predicted to be altered in multiple patients since they may point towards positively selected changes. Using 95^th^ percentile of reductions as a threshold to detect loops with significant decrease in loop probability, we identified fifteen loops with significant decrease in loop probability in at least 5% of patients (10/241 patients). There are fourteen genes inside these 15 loops. Among these fourteen genes, there are five known cancer genes: *RMI2, MYC, BCL2, LPP* and *SOCS1*. We used RNA-seq data from the 104 patients to identify differentially expressed genes among these fourteen genes. *BCL2* shows differential expression in the group of patients with decrease in loop probability vs. the other patients (Figure 4b). We did not find a significant difference in copy number of BCL2 between the two groups (p-value ~ 0.198), suggesting that changes in the strength of insulator loops could contribute to the increase in expression of *BCL2*.

## 3. Discussion

We developed a powerful method, DeepMILO, for modelling insulator loops and for predicting the effect of variants on these loops. DeepMILO has learned sequence features of insulator loops that are beyond the presence and orientation of CTCF motifs. It can identify insulator loops with high AUROC and performs significantly better than the model based on word2vec features and boosted trees. DeepMILO uses features learned by a CNN model and an RNN model. We show that RNN can perform comparably with CNN in learning DNA sequence patterns of insulator loop boundaries and that combining learned features of RNN and CNN models delivers a better model compared with individual models. Furthermore, we find small mutations of one base can result in significant impact on 3D chromatin loops and that our model is sensitive enough to predict the impact of such small mutations. Using DeepMILO, we identified constitutive insulator loops that change in multiple cancer patients and genes affected by these loops. Our model suggests a possible new mechanism for upregulation of the cancer driver gene BCL2 in malignant lymphoma via alteration of its 3D insulator loops. Our method is one of the first methods for predicting the effect of variants on 3D insulator loops. It can be extended for other 3D structures such as enhancer-promoter loops and TADs to identify the non-coding variants altering 3D chromatin structures. Identification of these variants together with altered 3D structures can provide insights into the mechanism of the aberrant expression of genes in cancer. Moreover, in future, the CNN and RNN models can be extended to use the attention mechanism ^27^ to allow them to focus more on important parts of input sequences to make them more robust.

## 4. Methods

### Data preparation

We used CTCF/cohesin chromatin loop data of the cell lines GM12878 and K562 to train and test our models. These loops were captured by cohesin (RAD21) ChIA-PET and their PET peaks were overlapped with CTCF ChIP-seq peaks ^5^. CTCF ChIP-seq data of GM12878 and K562 were obtained from ENCODE ^28^.

Boundaries of loops were normalized as following. Two boundaries *b*_1_ and *b*_2_ with lengths *l*_1_ and *l*_2_ are considered as equal and merged into one boundary if their overlap is larger than 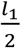 or 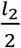. After merging equal boundaries, we obtained a set of normalized boundaries and loops were formed from these boundaries. Boundaries were then trimmed off or expanded to have a length of 4,000 bases (~ median length of boundaries captured by cohesin ChIA-PET) centered at their peaks. DNA sequences of boundaries were mutated with variants from corresponding cell lines. Variants of the cell lines GM12878 and K562 were obtained from the 1000 Genomes Project ^22 29^ and ^23^. All SNVs, indels, deletions, inversions, duplications and copy number variations that do not span the entire 4,000 bases of boundaries were used to obtain cell-line or patient-specific sequences. We modeled two alleles for each boundary. Sequence of an allele is converted into a one-hot encoding matrix of size [4000 x 5]. Due to the computationally intensive nature of LSTM, only 800 bases centered around the middle points of sequences were inputted into the RNN model.

Training data was derived from the cell line GM12878. There are ~19,000 boundaries and ~16,000 insulator loops in the cell line GM12878. Data from chromosomes 7 and 8 was held out for testing and data from chromosome 16 was used for validation. Additionally, data from chromosomes 7 and 8 of the cell line K562 CML was also used for testing. Negative boundary samples containing CTCF ChIP-seq peaks were derived from CTCF ChIP-seq signal and were centered around the middle points of peaks. Negative boundary samples containing CTCF motifs were calculated from location of CTCF motifs detected by FIMO ^30^ and they were also centered around CTCF motifs.

Positive and negative samples were balanced in training, validation and test sets. For each sequence, its complementary reverse sequence was also included to increase the amount of training data. Additionally, by training the model with complementary reverse sequences, the strand of sequences can be ignored when running the model. In preparation of the data to train the ‘boundary orientation model’, we removed 912 boundaries that can be considered as both left and right boundaries. Left and right boundaries were considered as negative and positive samples respectively.

### CNN, RNN and boundary model for learning features of boundaries

We developed a deep convolutional neural network (CNN) and a recurrent neural network (RNN) with bidirectional long short-term memory cells to learn DNA sequence patterns of boundary of insulator loops. Learned features from these two models were combined to build the boundary model to distinguish boundaries from non-boundaries.

DNA sequences of boundary and non-boundary regions were used to train the models. In training, validation and test sets, positive samples include true boundaries and negative samples consist of the three types of non-boundaries with a ratio of 50% : 30% : 20% for non-boundaries type 1, 2 and 3 respectively.

A DNA sequence was converted to a one-hot encoding matrix with *m* rows and 10 columns, where m is the length of the sequence and 10 columns corresponding to 5 bases A, C, G, T and N for two alleles. Modelling two alleles allows us to evaluate effect of heterozygous variants more accurately. The models were trained to output a number in the range of [0,1] that can be interpreted as boundary probability of a given DNA sequence. It is expected that probabilities of DNA sequences of boundary regions are closer to 1 and probabilities of DNA sequences of non-boundary regions are close to 0.

The CNN and RNN models were trained separately and their features were later combined for the boundary model. The same training and validation datasets were used to train the three models. The CNN model has two convolutional layers. The first layer of the network is a convolutional layer with 1024 filters of size [17 x 5]. Filters scan through input sequences and are applied to each allele separately. This layer is followed by a batch normalization ^31^, a leaky rectified linear and a dropout layer ^32^. The batch normalization layer stabilizes the output from the first layer before it goes through the leaky rectified linear layer and speeds up the optimization during training. The dropout layer prevents the network from overfitting. Our experiments found that a negative slope coefficient of 0.2 for the leaky rectified linear function and a dropout rate of 0.3 produced the best result. Following these layers is a second convolutional layer with 1024 filters of size [5 x 1]. This convolutional layer is supposed to combine features from the first convolutional layers to learn higher level features. Our experiments showed that this convolutional layer allowed us to achieve a better performance with less training time. However, adding more convolutional layers did not yield a clearly better performance while significantly increasing the complexity of the model. The second convolutional layer is followed by a batch normalization, a leaky rectified linear, a global max pooling and a dropout layer. The global max pooling layer is applied to alleles separately and is intended to capture if the input contains specific patterns learned by filters of convolutional layers. The output from these layers is then concatenated and inputted into two fully connected layers with 512 and 256 nodes respectively. The last layer of the model is a sigmoid activation node that outputs a probability of the input sequence as boundary of an insulator loop. We used a binary cross entropy loss function as objective function. And it was minimized using the RMSprop algorithm.

The RNN model consists of two stacked bidirectional LSTM layers (BLSTM) that can capture long-term dependency in long sequences. Bidirectional LSTM processes sequences in both directions, forward and backward directions, and therefore often captures the context better. Each BLSTM layer has 64 hidden units. A dropout layer with a dropout rate of 0.2 follows each layer to prevent overfitting. Output from BLSTM layers is inputted into a time distributed layer. Following the time distributed layer are fully connected layers with the same settings as in the CNN model.

To combine the learned features of the CNN and RNN models, their fully connected layers and output layers were stripped down after training and outputs from the remaining layers of the two models were concatenated and inputted into two new fully connected layers with 512 and 256 nodes as in the CNN and RNN model. The models were implemented in Keras (https://keras.io).

### Boundary orientation model to distinguish left and right boundaries

We constructed the boundary orientation model from learned features of the boundary model. The two fully connected layers of the boundary model were replaced by two new fully connected layers of 512 and 256 nodes respectively. We then trained this model to distinguish left and right boundaries of insulator loops. Left and right boundaries were considered as negative and positive samples respectively. The model is expected to produce values close to 0 for left boundaries and values close to 1 for right boundaries.

### Loop model for learning sequence patterns of insulator loops

We built DeepMILO by combining learned features of the boundary and boundary orientation models as shown in (Figure 3b). The output layer of the boundary model was removed and the two fully connected layers were replaced by two new layers with the same settings to make a new sub-model. Then, the output from this new sub-model was concatenated with outputs from the boundary and boundary orientation model. The loop model uses sequence features of boundaries and predicted outcomes from the boundary model and the boundary orientation model for its prediction. In training, validation and test sets, positive samples are true insulator loop and negative samples consists of non-loop type 1, 2, 3, 4 and 5 with a ratio of 50% : 10% : 10% : 20% : 10% respectively.

### Implementation of a classifier using word2Vec and boosted trees

Following the CTCF-MP model in ^17^ we implemented a classifier using word2Vec and boosted trees to identify loops using DNA sequence only. CTCF-MP utilizes both genetic and epigenetic features (including CTCF ChIP-seq) to make prediction, however, using only DNA sequence features from word2vec, it predicted CTCF ChIA-PET loops with AUROC of 0.796 ^17^. We implemented a similar model that takes DNA sequence features only as input to compare with DeepMILO. Boundaries were represented by DNA sequences of size 520 bases centered around their middle points. Our implementation achieved a comparable average AUROC of 0.794 in a 10-fold cross-validation using the same dataset by CTCF-MP ^17^. We then trained the classifier with the same training datasets used for DeepMILO. And the classifier was tested with the five test sets from GM12878 to compare with DeepMILO. To improve performance of this model, we also tried to increase the boundary size to allow the model to capture more context of the data but found that the model generally performed worse with longer boundary sizes.

### Program availability

The source code and models are available at: https://github.com/tuantrieu/DeepMILO

---

**Figure S 1.**
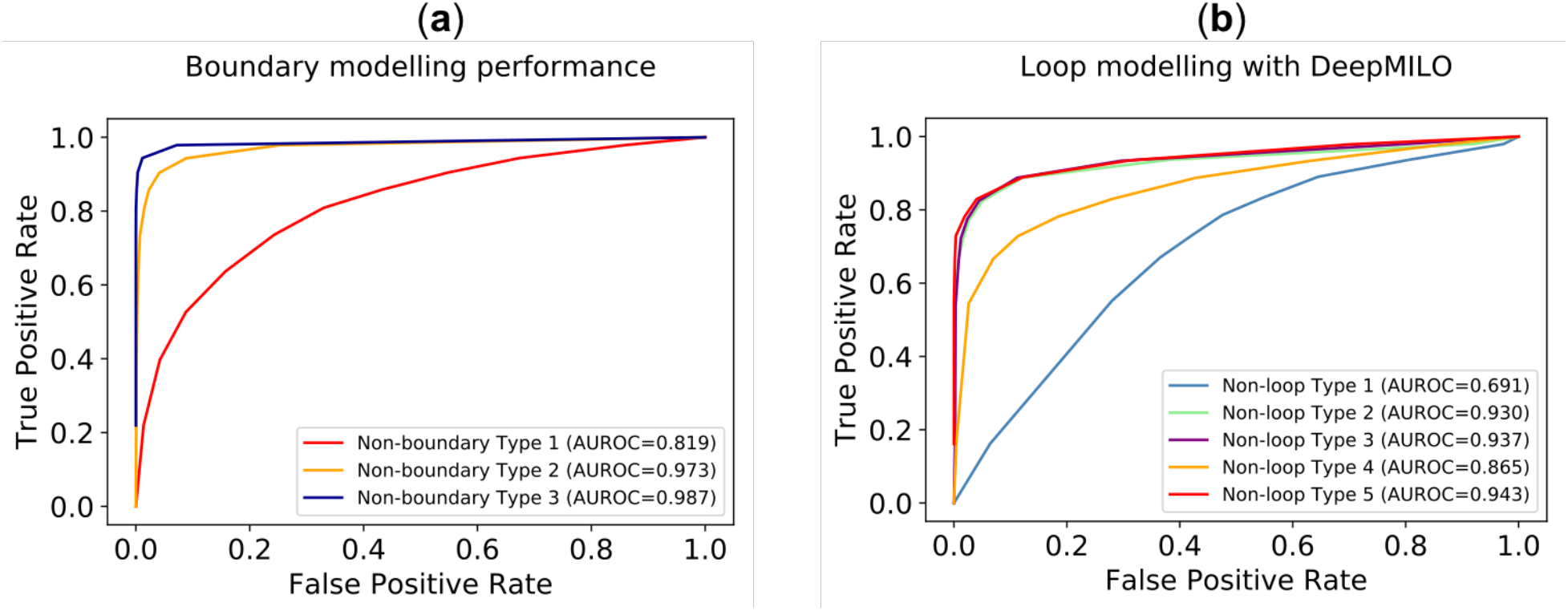
(**a**) Boundary modelling performance on test sets from K562 (**b**) Performance of loop modelling using deep learning on test sets from K562

**Figure S 2.**
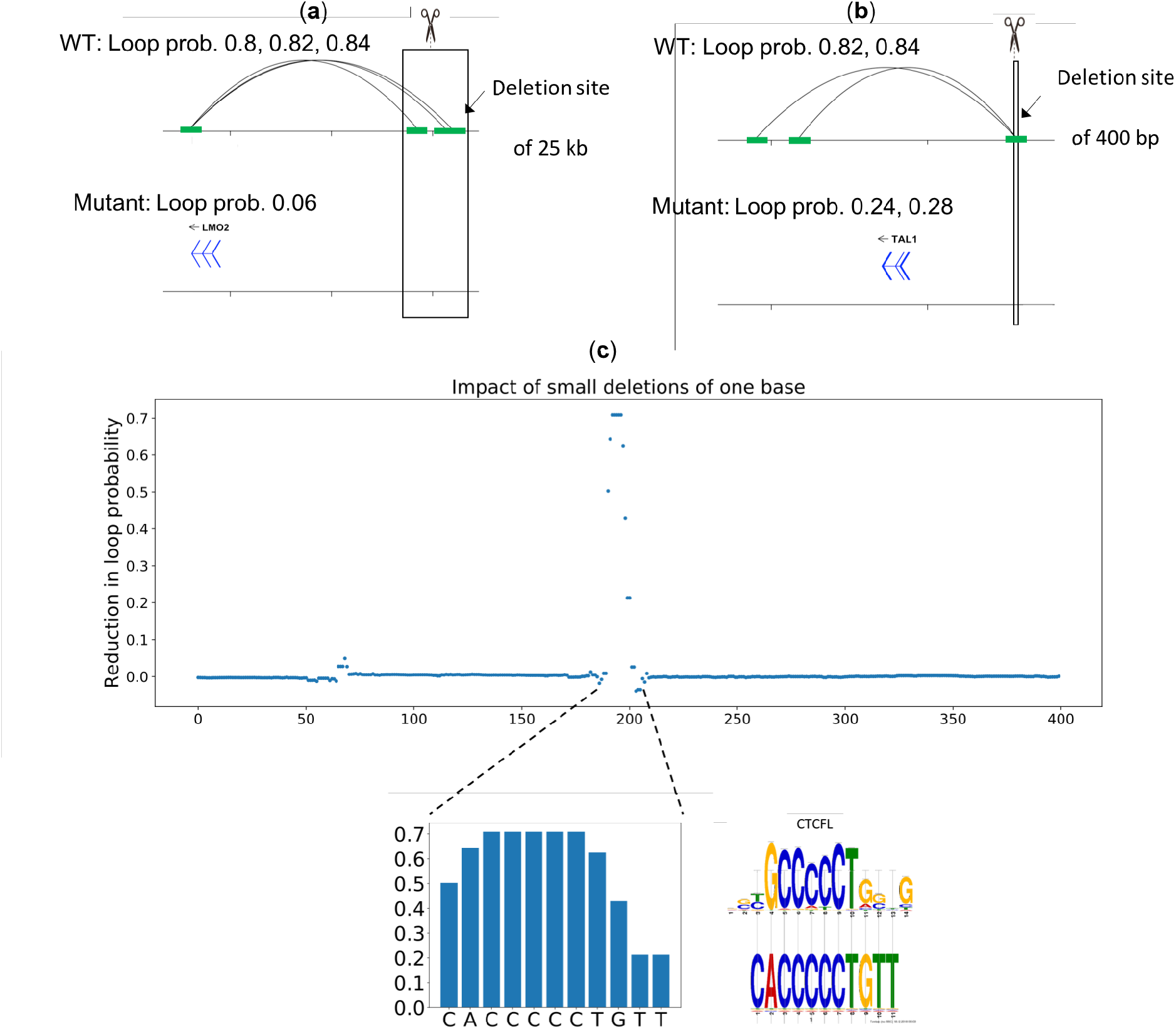
Validation of the loop model with known deletions that disrupt insulator loops. (**a**) Insulator loops cover LMO2 gene. The deletion decreased loop probabilities from ~0.8 to 0.6 (**b**) Insulator loops cover TAL1 gene. Loop probabilities decreased from ~0.82 to 0.24 because of the deletion. (**c**) Impact of 400 small deletions on a loop containing TAL1.

